# CRISPR-DIPOFF: An Interpretable Deep Learning Approach for CRISPR Cas-9 Off-Target Prediction

**DOI:** 10.1101/2023.08.05.552139

**Authors:** Md. Toufikuzzaman, Md. Abul Hassan Samee, M Sohel Rahman

## Abstract

CRISPR Cas-9 is a groundbreaking gene-editing tool that harnesses bacterial defense systems to alter DNA sequences accurately. This innovative technology holds vast promise in multiple domains like biotechnology, agriculture, and medicine. However, such power does not come without its own peril, and one such issue is the potential for unintended modifications (Off-Target), which highlights the need for accurate prediction and mitigation strategies. Though previous studies have demonstrated improvement in Off-Target prediction capability with the application of deep learning, they often struggle with the precision-recall trade-off, limiting their effectiveness and do not provide proper interpretation of the complex decision-making process of their models. To address these limitations, we have thoroughly explored deep learning networks, particularly the recurrent neural network (RNN) and transformer based models, leveraging their established success in handling sequence data. Furthermore, we have employed genetic algorithm for hyperparameter tuning to optimize these models’ performance. The results from our experiments demonstrate significant performance improvement compared to the current state-of-the-art in Off-Target prediction, highlighting the efficacy of our approach. Furthermore, leveraging the power of the integrated gradient method, we make an effort to interpret our models resulting in a detailed analysis and understanding of the underlying factors that contribute to Off-Target predictions, in particular the presence of two sub-regions in the seed region of sgRNA which extends the established biological hypothesis of Off-Target effects. To the best of our knowledge, our model can be considered as the first model combining high efficacy, interpretability, and a desirable balance between precision and recall.

## Introduction

CRISPR Cas-9 stands for Clustered Regularly Interspaced Short Palindromic Repeats and CRISPR-associated enzyme Cas-9. It utilizes a bacterial defense mechanism to precisely cut DNA sequences which has been repurposed by scientists to cause a double strand break at any portion of the DNA by providing the Cas-9 enzyme with the exact sequence of that DNA portion [24]. After the break, DNA’s natural healing mechanism is triggered and due to mutation, the repair region’s biological function gets disrupted. Thus, CRISPR Cas-9 system can essentially be exploited to knock off specific genes in an organism’s genome. This repair mechanism can be controlled by providing a template to replace the existing DNA sequence [39, 57] and thus the gene gets edited.

The applications of CRISPR are beyond imagination. At the very least, CRISPR can be used to map the functionality of genes in different organisms. CRISPR can revolutionize the agriculture sector by designing disease and weed resistant high yielding crops [62]. Another application of CRISPR is preventing and curing genetic diseases. Sickle Cell Anemia is such a disease which affects hemoglobin due to an unintended mutation in *β*-globin gene. Once the only treatment for this disease was bone marrow transplant. By knocking off the faulty gene and correcting it with a controlled repair mechanism CRISPR has opened up a new approach of treating sickle cell anemia [4] and its human clinical trial has also started [16]. Scientists have already provided theoretical proof of concept on how gene drive on mosquitoes using CRISPR could wipe out the deadly disease malaria [21]. However, like any other groundbreaking advancement, CRISPR comes with its own set of challenges, particularly in the form of Off-Target effects. Identifying these Off-Targets through traditional laboratory validation methods can be costly and time-consuming. This presents a valuable opportunity for computational methods to offer a rapid, reliable, and cost-effective solution to detect and predict Off-Target effects in gene editing.

The detection of potential Off-Target effects is very crucial for the successful application of the CRISPR Cas-9 system. In recent years, computational approaches, specially machine learning based ones, have contributed to a great extent in fields like genomic sequence analysis, drug discovery, motif finding etc. [40]. Following the same trend, computational approaches have been used to predict the Off-Target effects too. The early detection methods relied on score based approaches [23, 50, 49, 15] primarily focusing on identifying Off-Target effects based on the presence and positions of mismatches between the target site and the guide RNA assigning scores based on the mismatches. These methods do not consider the complex relationship among the sequences and they may suffer from experimental variations and limitations [33].

Traditional machine learning approaches for Off-Target prediction typically involve the use of manually engineered features. These features can include sequence-based characteristics, such as mismatch patterns or nucleotide composition as well as contextual information surrounding the target site. The machine learning algorithms are trained on labeled datasets to learn the relationship between the features and the Off-Target activity. Abadi et. al. [1] aggregated datsets from three different sources: GUIDE-Seq [52], HTGTS [30], and BLESS [18], and trained a random forest regression model. Peng et. al. [42] trained a number of SVM (Support Vector Machine) models where number of positive samples and negative samples were maintained the same. The models were ensembled using the average prediction probability of all the models. Chen et. al. [8] performed their experiments on their CRISPEY dataset and applied SMOTE (Synthetic Minority Over-Sampling Technique) [6] to generate synthetic positive sample and trained Logistic Regression, SVM, Random Forest and Neural Network models. Zhang et. al. [59] used the scores of score-based methods along with biological properties as features for training an Adaboost [17] model.

Over the time, the dataset for Off-Target prediction has grown significantly and recently most of the works have focused on deep learning for predicting Off-Target site as traditional machine learning models do not scale well for large datasets [47]. Lin et. al. [33] presented a CNN (Convolutional Neural Network) based model CNN Std trained on CRISPOR [20] dataset. They encoded the sgRNA and targeted DNA sequences with the one hot encoding method and superposed (logically ORed) those encodings. In a similar study, Charlier et. al. [5] concatenated the one hot encodings of the sequences instead of superposing those and their FNN (Feedforwad Neural Network) model outperformed CNN Std. Chuai et al. [10] produced their own dataset (DeepCRISPR) which has been used by most of the later works as a benchmark dataset. The authors pretrained a DCDNN (Deep Convolutionary Denoising Neural Network) based autoendcoder using the whole human genome sequence containing almost 0.68 billion unlabeled samples. They used epigenetic features along with the sequence data. In the downstream of the autoencoder, there were CNN and fully connected layers before the output layer.

Natural Language Processing (NLP) based techniques have also been employed for Off-Target prediction. For example, Liu et. al. [35] applied transformers [53] for Off-Target prediction in their proposed tool AttnToMismtach CNN. The model was a combination of sequence and position embedding, transformers and CNN. Lin et. al. [34] presented a prediction tool named CRISPR Net which uses a 7 channel encoding scheme, where in addition to the 4 channels of CNN std [33] there is one channel for insertion or deletion type mismatch detection and two channels for direction of mismatch. Similar approach was adopted by the tool CRISPR-IP [60]. Liu et. al. [36] proposed CnnCrispr where they used the dataset of DeepCRISPR [10] as benchmark. They introduced GloVe (Global vectors for word representation) embedding [43] for the input sequences. They reported significant improvement on AUPRC (Area Under Precision Recall Curve) score over the existing models.

Vinodkumar et. al. [54] applied GCN (Graph Convolutions Network) [29] and employed a link prediction method to predict the presence of links between the single guide RNA (sgRNA) and target sequences that generate Off-Target effects. Chen et. al. [7] pretrained a BERT [14] model specifically targeting the Off-Target prediction task. They pretrained their BERT model using the similar pretraining data of DeepCRISPR. Instead of fine tuning the pretrained model, they later used the BERT model’s embedding as features and along with other engineered features trained a LightGBM model [25].

Though the application of deep leaning has improved the accuracy of Off-Target prediction, the datasets of Off-Target prediction are highly imbalanced and thus most of the models suffer from precision-recall trade-off. Another shortcoming of the existing studies is the lack of focus on interpreting the models. A few traditional machine learning models applied feature rankings [1, 44, 38] for On-Target and Off-Target prediction whereas most of the deep learning models do not provide any interpretation at all. A few deep learning models provide brief interpretation with saliency map [10] or input perturbation based feature ranking [35] but they do not explore the complex decision making process with a in-depth analysis. Due to the sensitiveness in the application of CRISPR in real life, it is absolutely necessary to understand the inner workings and decision making process of the prediction models.

In this study, we have attempted to address existing issues in Off-Target prediction. This paper makes the following key contributions.

We present CRISPR-DIPOFF suite, a set of interpretable deep learning models capable of accurately predicting Off-Target sites using only sequence data (Section 2).

- In parallel, we also present a framework utilizing genetic algorithms that enables the optimization of hyperparameters for deep learning models (Section 3.1). CRISPR-DIPOFF utilizes this framework to achieve improved performance and generalization.
- CRISPR-DIPOFF has been effectively interpreted using integrated gradients [51], establishing links with established biological hypotheses (Section 4). Specially our study has identified two possible sub-regions in the seed region one of which shows positive correlations with Off-Target effects.
- Finally, to facilitate replication and extension of the proposed algorithms, a modular implementation has been developed and has been made available ^1^.

## Methods

### Dataset

In this study, we utilize two distinct datasets. Each dataset serves a specific purpose and contributes to the overall objective of our work. The datasets are briefly described below.

#### DeepCRISPR Off-Target Dataset

The dataset used in the DeepCRISPR [10] study, serves as a foundational resource for our analysis. It comprises a comprehensive collection of single guide RNA (sgRNA) sequences along with their corresponding potential Off-Target sites and the dataset was labeled using various whole genome Off-Target detection methods [52, 27, 26, 18]. Since publication, this dataset has been widely used in different studies of CRISPR Cas-9 Off-Target prediction. The dataset contains samples from two cell lines: k562 (12 sgRNAs) and hek293 (18 sgRNAs) [23, 9]. For these 30 sgRNAs, the authors collected 153233 loci across the whole genome using a tool named Bowtie 2 [32]. They allowed up to six mismatches between sgRNA and the target DNA. The dataset is highly imbalanced. It has roughly one positive (label 1) Off-Target sample for each 230 negative (label 0) negative samples.

For our experiment we split the dataset into three subsets: training (60%), validation (20%), and test (20%). The purpose of these subsets is to train the model, tune hyperparameters, and evaluate its performance, respectively.

#### Human Genome Sequence (GrCh38)

we further employed a pretraining approach using a transformer-based ELECTRA [12] model. The objective was to train the model on the vast and comprehensive human genome sequence data. To accomplish this, we utilized the GRCh38 (Genome Reference Consortium Human Build 38) as the dataset for pretraining which is a comprehensive assembly of the human genome with improvements over previous version, GRCh37 [46]. It includes the autosomes (chromosomes 1 to 22) as well as the sex chromosomes (X and Y) and mitochondrial chromosomes. We collected this dataset from the Ensembl [58] project’s repository ^2^.

### Performance Metrics

In order to evaluate the performance of our models and assess their efficacy in predicting Off-Target effects, we have employed various performance metrics like accuracy, precision, recall, F-1 score, AUROC (Area Under the Receiver Operating Characteristics) and AUPRC (Area Under the Precision-Recall Curve). Most of the previous studies reported and compared their results in terms of AUROC score. But AUPRC is often considered a better measure for imbalanced datasets compared to AUROC [13, 35]. In imbalanced datasets, where the number of negative instances (non Off-Target sites) far exceeds the number of positive instances (Off-Target sites), the AUROC can be misleading as it calculates the performance based on the true positive rate (recall) and false positive rate, regardless of the class distribution. As a result, if the majority class dominates the dataset, the AUROC can appear high even if the model performs poorly on the minority class. On the other hand, the AUPRC takes into account the precision and recall, which are more informative for imbalanced datasets. It focuses on the ability of a model to correctly identify positive instances (Off-Target sites) while maintaining a low false positive rate. Hence in our study, we have used AUPRC as the primary metric for comparison.

### Recurrent Neural Network-Based Approach

The Off-Target prediction task involves analyzing two input sequences: sgRNA (single guide RNA) and target DNA. The objective is to determine the likelihood of potential Off-Target sites. To achieve this objective, we were motivated to explore the Recurrent Neural Networks (RNNs) and their variants. RNNs have gained significant popularity and success in sequence prediction tasks across various domains, such as natural language processing, speech recognition, time series analysis, and genomics. While previous works have employed LSTM (Long Short-Term Memory) [35, 59], it has typically been used as a component within a larger model. We explored different variants of RNN, like vanilla RNN, LSTM and GRU (Gated Recurrent Unit) and their different compositions such as, Bi-directional RNN and Stacked RNN.

#### Data Preprocessing

First we preprocessed the raw sgRNA and target DNA sequences using two one hot encoding methods (4-channel and 5-channel) as shown in Figure 1. We followed the same approach used by CNN Std [33]. For, 4-channel encoding, we converted both sgRNA and DNA sequence to one hot encoded matrix independently. There are four channels corresponding to four possible nucleotides. This process produced a 4 *×* 23 sized binary matrix for a 23-mer sequence. Two such matrices, one for sgRNA and one for target DNA, were superposed using logical OR operation to produce the final encoding. We added an additional directional channel in 5-channel encoding method. In our encoding scheme, the precedence of encoding is ‘A’,’T’, ‘C’ and ‘G’. In case of a mismatch, if the 1 corresponding to the higher precedence nucleotide originates from the target DNA sequence, the direction channel is set to ‘1’.

**Fig. 1.**
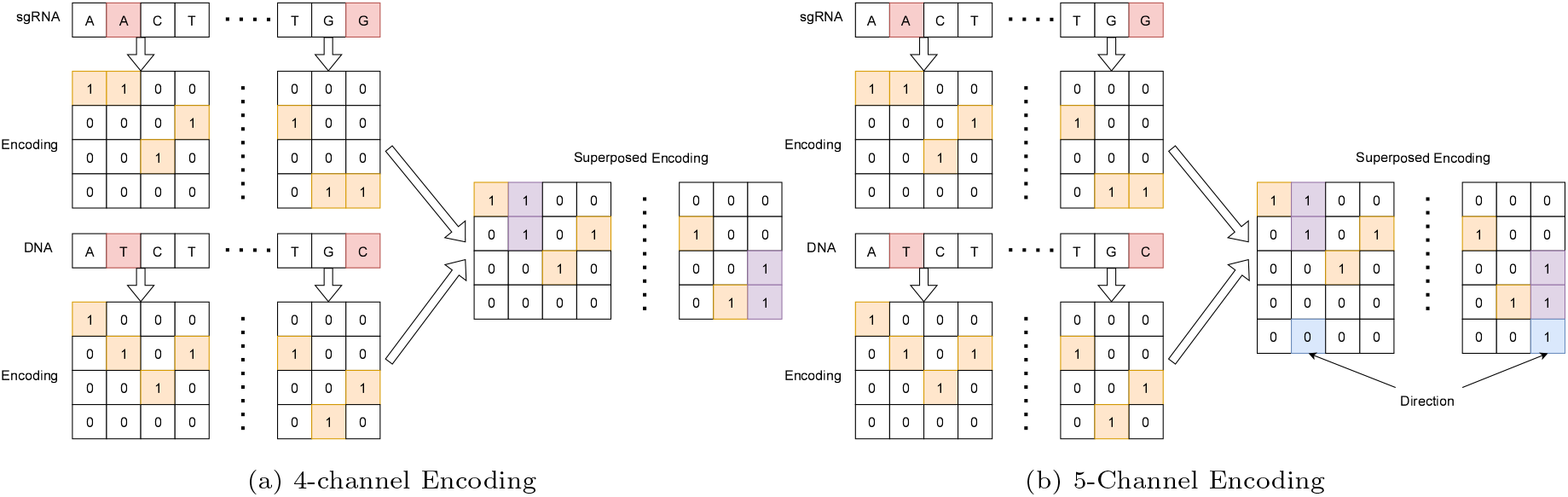
Example of One Hot Encoding. (a) The sgRNA and DNA sequence are encoded with 4-channels independently and then superposed using logical OR operation. (b) An additional of channel has been added for direction.

#### Generic Model Design

We designed a generic parameterized RNN model which would ease the process of hyperparameter tuning. A simplified block diagram of the model is shown in Figure 2. The model’s input would be the 4/5-channel matrix generated earlier from the raw sequences. The encoded sequences would then be fed to a RNN/LSTM/GRU layer. Based on the parameter there could be one or two recurrent layers and each layer could be bidirectional or unidirectional. The output from the RNN/LSTM/GRU layer is passed through a series of fully connected hidden layers each of which gradually halves the size of its previous layer. Finally, there is an output layer. The output layer’s activation function is logit and hidden layer’s activation function is ReLU. We used CrossEntropy as the loss function and Adam (Adaptive moment estimation) [28] as the optimizer. All the hidden layers were followed by the ReLU activation layer and a dropout layer to tackle overfitting during training.

**Fig. 2.**
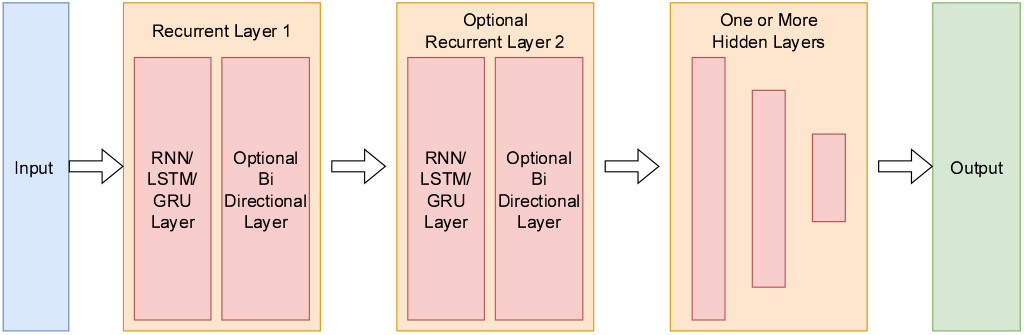
Block Diagram of Generic RNN Based Model. The encoded input matrix passes through one or two uni or bi-directional Recurrent layers, followed by a series of hidden layers. There are ReLU activation and dropout layer in between the hidden layers.

#### Model Selection with Genetic Algorithms

In order to find the best set of hyperparameters for our model we performed a structured model selection process using genetic algorithms which are optimization algorithms inspired by the process of natural evolution. Most of the previous works performed model selection only in a limited scale and that too mostly based on trial and error or adding or removing different components of the model [33, 36]. We selected the eight hyperparameters shown in Table 1 to be tuned for our generic model. Many of the parameters take on continuous values. But due to our limitation of computational resources we prepared a list of possible values for each parameter to be optimized. The values were chosen from the range which is generally used in deep learning models and suggested by different empirical studies [55].

**Table 1.**
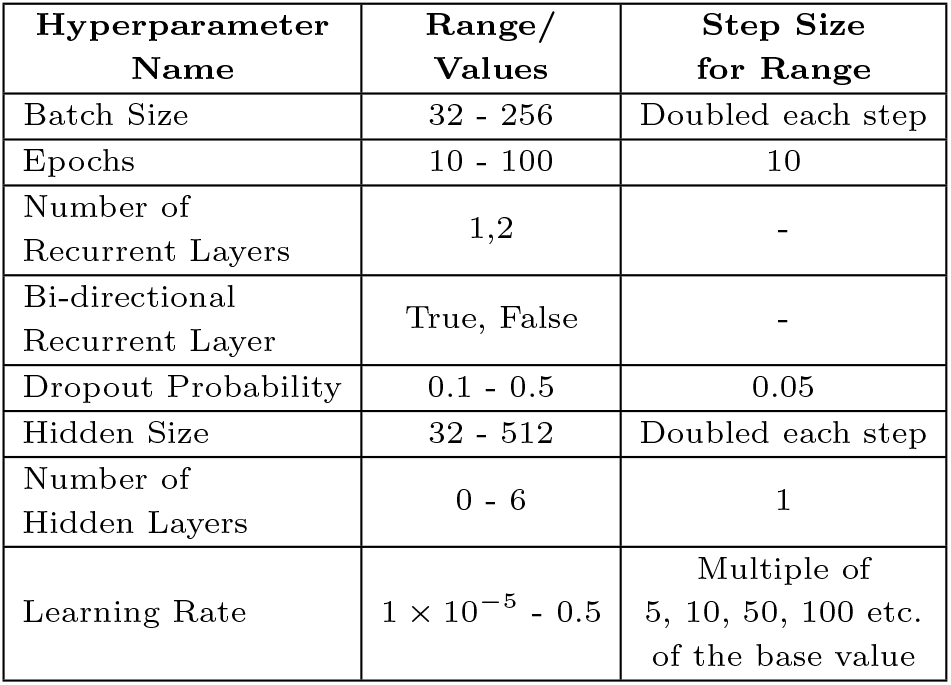
List of hyperparameters and their possible values used in our study.

Among the hyperparamters, the possible values for the number of hidden layers are dependent on the hidden size of the RNN unit. Even without including this hyperparameter, there would be 72,000 possible combination of the hyperparameters. Training each model on our computational environment took around 10 to 30 minutes based on the selected hyperparameters. In our case, if we were to extensively search through all possible combination of hyperparameters for a single type of recurrent network (RNN/LSTM/GRU), it would take about 1000 days! Hence we choose to apply genetic algorithm to intelligently explore the search space. The steps of genetic algorithm are discussed below.

- **Initial Set of Model Parameters**: First we prepared a set of hyperparameters by randomly selecting from the possible values for each of the initial models (population).
- **Model Training**: This is the bottleneck step of the whole process. Using the set of hyperparameters, each model is trained using the training set (60% of the whole dataset).
- **Fitness Assessment**: After the training process, each model’s performance is evaluated using the validation set (20% of the whole dataset). AUPRC was chosen as the objective function. We ranked the models of each generation based on the AUPRC score.
- **Model Selection**: We explored with two variants of genetic algorithm. In the plain genetic algorithm variant, new set of model hyperparameters are generated using the hyperparameters of the last generation and the previous generation’s hyperparameters are discarded. This actually causes the already found “useful” models to be lost in the selection process. In another variant of genetic algorithm, popularly known as Elitist Genetic Algorithm, we selected a certain number of models (Elites) with higher fitness score which were passed on to the next generation along with the new set of models.
- **Crossover of Model Parameters**: In order to generate new set of hyperparameters, we first selected models by pairs through tournament selection where we first choose a model randomly and for the specified tournament size, we continued selecting models randomly and returned the model with highest fitness. Then we performed crossover between the hyperparameters of the models which is the process of iteratively going through the hyperparameters and for each hyperparameter, with a certain probability, values were swapped between the models. This pairwise crossover process continued until the required number of new hyperparameter sets had been found.
- **Mutation of Model Parameters**: In this process, each set of hyperparameters from crossover are considered one by one. A few hyperparameters with a specified probability are subjected to mutation, which actually means that their values are substituted with randomly selected values from the possible set of values.
- **New Set of Model Parameters**: After the mutation process, we were left with a new set of hyperparameters. Model with these parameters were trained and the steps continued iteratively until a certain number of generations have passed. At the end, the process returned the best performing model’s hyperparameters.

In our study, the population, elite and tournament size were 20, 4 and 2 respectively. The crossover and mutation probabilities were 0.3 and 0.2. The genetic algorithm was run for 20 generations. Each of RNN, LSTM and GRU models works a bit differently so it was not conducive to perform crossover among different types of recurrent networks. Hence we run genetic algorithm and its elitist variant for each of the recurrent model types independently. The best RNN, LSTM and GRU models found from these experiments were trained with the combination of training and validation set and evaluated on the test set for comparison with baselines.

### Transformer-Based approaches

In recent years, the development of transformer-based models, such as BERT (Bidirectional Encoder Representations from Transformers)[14] and ELECTRA (Efficiently Learning an Encoder that Classifies Token Replacements Accurately) [12], has revolutionized natural language processing tasks. One key aspect that has contributed to the success of these models is pretraining which involves training the models on large-scale datasets, allowing them to learn general language representations. We designed an ELECTRA based pipeline to utilize large pretrained model for Off-Target prediction. ELECTRA is considered an improvement over BERT due to its adversarial training approach, which leads to better model generalization. It is computationally more efficient, requiring fewer parameters and less pretraining time. ELECTRA also exhibits improved robustness and stability, making it a preferred choice for natural language processing tasks [12].

#### Data Preprocessing

We followed a data processing approach similar to that used in DeepCRISPR’s [10] pretraining. But we used GRCh38 instead of GRCh37 as the data source for human genome sequence. We used the Cas-OFFinder [3] tool to find sample pairs that allowed up to 6 nucleotide mismatches where the first sequence (sgRNA) ends with ‘NGG’. This process resulted in a pretraining dataset comprising approximately 317 million samples. In our study, we explored two approaches to construct the vocabulary: overlapping tri-mer and byte pair encoding (BPE) [19]. Both the approaches have five special tokens: [UNK], [CLS], [SEP], [MASK] and [PAD] representing unknown, classification, separator, mask and padding tokens respectively. Overlapping tri-mer involves dividing the sequences into overlapping three-nucleotide segments. Considering four possible nucleotides, there are 4 *×* 4*×*4 = 64 possible tri-mer tokens. Along with the special tokens there are 69 tokens in the vocabulary generated by overlapping tri-mer encoding. A 23-mer sequence would be converted to a sequence of 21 tokens in this approach. The other approach, unigram Byte Pair Encoding (BPE), is a subword tokenization technique which involves iteratively merging the most frequent character pairs in a corpus to create a vocabulary of subword units. We trained the BPE algorithm on the pretraining dataset and it produced a total of 355 tokens including the 5 special tokens.

After forming the vocabulary we preprocessed the pretraining samples. Each sequence pairs were first tokenized. The tokens of two sequences were separated by a [SEP] token and a [CLS] token was inserted before the first token of sgRNA. Another [SEP] token was added at the end of the target DNA tokens. Maximum sequence length was set to 48 and the remaining positions after the input tokens were filled with [PAD] tokens. Input mask was generated to identify different sequences (0 for sgRNA and 1 for potential target DNA).

Attention mask differentiated the actual input tokens from the [PAD] tokens.

#### Pretraining the ELECTRA model

In the process of pretraining ELECTRA, a subset of approximately 15% of the samples was selected for replacement with a [MASK] token. This masked portion of the data was then passed through a generator, which substituted the masked tokens with generated tokens. The discriminator’s task was to determine whether the tokens had been replaced by the generator or not. To accommodate limited resources, we conducted pretraining using two relatively smaller versions of the ELECTRA model, namely, Tiny and Small models. The training process consisted of 100,000 steps with a batch size of 128. Towards the end of training, the loss function stabilized over the last few thousand steps. The two smaller versions of the ELECTRA model employed the parameter configurations shown in Table 2. We kept all other parameters consistent with the official ELECTRA code repository. The tiny model underwent training for approximately 7 days, while the small model was trained for 10 days. Both models were trained separately using tri-mer and byte pair encoded tokens.

**Table 2.** Parameters for ELECTRA Tiny and Small Model.

| Parameter | Tiny Model | Small Model |
| --- | --- | --- |
| Embedding Size | 128 | 512 |
| Hidden Size | 256 | 512 |
| Intermediate Size | 512 | 1024 |
| Encoder Layers | 6 | 8 |
| Attention Heads | 6 | 8 |

#### Finetuning ELECTRA Model for Off-Target Prediction

Following pretraining, the ELECTRA model was further finetuned on a specific downstream task, where it was trained on the labeled DeepCRISPR dataset. During finetuning, the model’s parameters were adjusted to optimize its performance on the task’s objective using supervised learning. We conducted experiments with different numbers of layers (ranging from 1 to 3) attached to the output of the [CLS] token of the ELECTRA model and finetuned them. Models pretrained on both tri-mer and byte pair encoded tokens were finetuned separately. Initially, during the finetuning process, the model faced difficulties in learning. It exhibited a tendency to predict all samples as either 1 or 0. To overcome this, we employed a training technique called gradual unfreezing, as described by Howard et. al. [22]. This involved initially freezing the weights of the ELECTRA model and updating only the output layer weights for a few iterations. Subsequently, we gradually unfroze the weights of each ELECTRA layer, starting from the layer closest to the output layer in a bottom-up manner. While this approach led to gradual improvement in the performance of the Tiny model, the Small model still exhibited a tendency to predict all samples as either 0 or 1. As a result, we have focused on comparing the results obtained using the Tiny model.

## Code, Environment and Availability

All the deep learning models were designed using the PyTorch deep learning library [41]. The recurrent neural network based experiments including the hyperparameter tuning with genetic algorithm was performed on an Nvidia RTX3090 GPU with 24GB VRAM. The ELECTRA model was pretrained and finetuned on an Nvidia Titan X GPU with 12 GB of VRAM. We used PyTorch Captum [31] library for model interpretation which provides the implementation of integrated gradients. The dataset and source codes are available at https://github.com/tzpranto/CRISPR-DIPOFF.

## Results

### Model Selection

In order to perform model selection, we ran plain genetic algorithm and elitist genetic algorithm (EGA) separately for 4-channel and 5-channel encoding scheme. Including the initial population, we trained a total of 420 models out of 72,000 (or even more) possible models in each experiment. The same experiment was run independently for RNN, LSTM and GRU type recurrent networks. We ranked and selected the models based on their AUPRC score on the validation set across all the experiments. The best RNN, LSTM and GRU model’s hyperparameters along with their performance on validation and test set are shown in Table 3. The detailed results of the individual experiments are presented in Appendix A.

**Table 3.** Hyperparamters of best RNN, LSTM and GRU models along with their AUPRC score on validation and test.

| Model Type | Hidden Size | LSTM Layers | Bi LSTM | Hidden Layers | Dropout Probability | Batch Size | Epochs | Learning Rate | AUPRC on Validation Set | AUPRC on Test Set |
| --- | --- | --- | --- | --- | --- | --- | --- | --- | --- | --- |
| RNN | 256 | 2 | TRUE | 0 | 0.4 | 256 | 60 | 0.00100 | **0.7643** | 0.5711 |
| LSTM | 512 | 1 | TRUE | 2 | 0.4 | 64 | 50 | 0.00010 | 0.7403 | **0.7208** |
| GRU | 128 | 2 | TRUE | 0 | 0.1 | 64 | 30 | 0.00050 | 0.7427 | 0.6859 |

According to the results, all the models have performed quite well on the validation set in terms of the AUPRC score. We observed that most of the models from plain genetic algorithm were already found on iteration 0, 1 or 2. This indicates that the best performing models from initial generation were not preserved in the subsequent generations and some useful parameter values might have got lost. This effect was reduced using the elitist version of the genetic algorithm which had produced the best performing RNN and LSTM models. Surprisingly, in both version of genetic algorithms, the performance of 5-channel encoded data was worse. This observation suggests that the mismatch itself might be the primary factor influencing Off-Target prediction, and incorporating the direction of the mismatch as an engineered feature might have introduced unnecessary complexity or bias to the models.

The selected best models were trained on the combined training and validation set and their performances were measured on the test set for further comparison with previous studies. The AUPRC scores on the test set have decreased compared to the AUPRC score on validation set and it is more prominent for the RNN model (rightmost two columns of Table 3). In general LSTM and GRU can generally store long term dependency and capture complex pattern compared to RNN [11].

### Finetuned ELECTRA Models

During the finetuning phase of the ELECTRA models, we experimented with different number of additional layers, ranging from 1 to 3, including the output layer and employed gradual unfreezing [22]. We conducted separate finetuning experiments on the Tiny models that were pretrained using both Tri-Mer Encoded (TME) and Byte Pair Encoded (BPE) input tokens. The detailed results of these experiments, have been reported in Appendix B. Notably, we observed a significant performance advantage for the TME models compared to the BPE models. Among the TME models, the one that yielded the best results incorporated two additional layers, an additional hidden layer and an output layer, following the ELECTRA encoder layers. The results of this model and is shown in Table 4.

**Table 4.** Results of best finetuned ELECTRA model. This model was pretrained with Tri-Mer Encoded (TME) input tokens and it included two additional layer for classification. The result shows how the various performance metrics improved over the gradual unfreezing of ELECTRA layers.

| Model Type | Unfrozen Layers | Trainable ELECTRA Parameters | Accuracy | Precision | Recall | F1-Score | AUROC | AUPRC |
| --- | --- | --- | --- | --- | --- | --- | --- | --- |
| TME with 2 FCL | None | 0 | 0.8078 | 0.0197 | 0.9008 | 0.0385 | 0.9264 | 0.0657 |
|  | Layer 6 | 527104 | 0.8077 | 0.0208 | 0.9542 | 0.0407 | 0.9369 | 0.1009 |
|  | Layer 5 to 6 | 1054208 | 0.8244 | 0.0229 | 0.9618 | 0.0447 | 0.9479 | 0.1078 |
|  | Layer 4 to 6 | 1581312 | 0.8243 | 0.0231 | **0.9695** | 0.0450 | 0.9572 | 0.1161 |
|  | Layer 3 to 6 | 2108416 | 0.9380 | 0.0576 | 0.8779 | 0.1080 | 0.9752 | 0.2846 |
|  | Layer 2 to 6 | 2635520 | 0.9772 | 0.1271 | 0.7405 | 0.2170 | **0.9775** | 0.3596 |
|  | Layer 1 to 6 | 3162624 | 0.9857 | 0.1848 | 0.6870 | 0.2913 | 0.9742 | 0.3951 |
|  | All Layers | 3194112 | **0.9898** | **0.2378** | 0.6336 | **0.3458** | 0.9752 | **0.4613** |

Though the overall result does not outperform the RNN based models, we observe that gradual unfreezing improves the performance of the models by solving the problem of catastrophic forgetting [22] which causes the model to forget the pretrained general knowledge if all model weights are updated at once. Figure 3 shows the effect of gradual unfreezing on performance metrics for the best performing ELECTRA model. It shows that most of the metrics have improved gradually except recall. The initial model had the tendency to predict a lot of Off-Targets which affected the precision of the model. As the model gradually stroke balance between precision and recall, precision gradually increased and recall gradually decreased, with overall improvements in F1-Score and AUPRC.

**Fig. 3.**
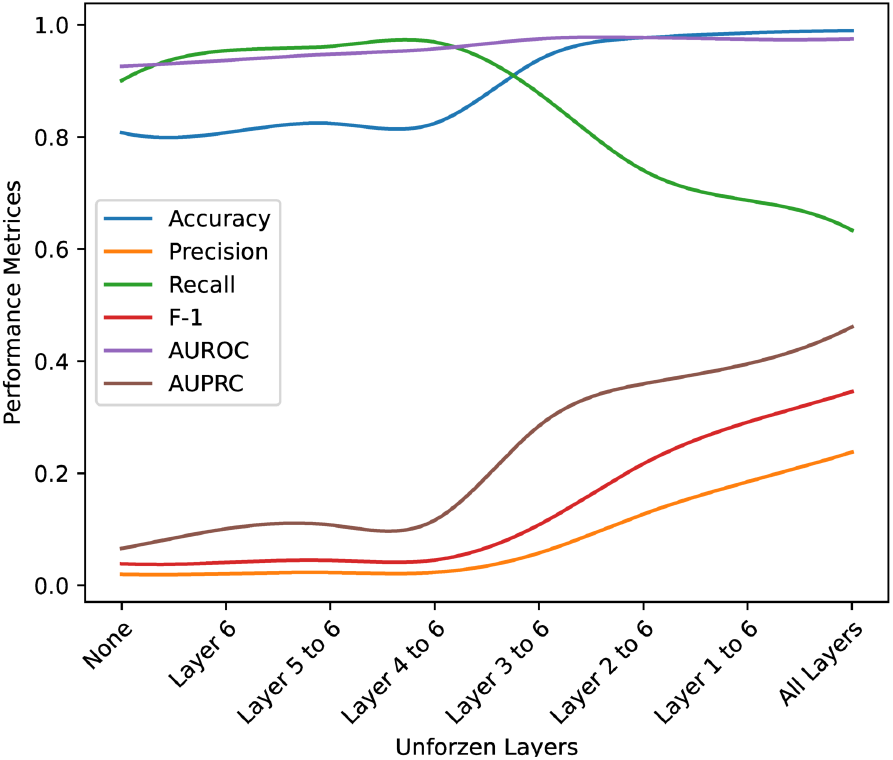
Effect of gradual unfreezing on different performance metrics. All the performance metrics have increased gradually when the parameters of the ELECTRA layers were unfreezed for finetuning one by one. The only exception was recall as it decreased with the increase of precision.

### Comparison with Previous Studies

We retrained the deep learning models from previous studies whose source code is publicly available among the models discussed in Section 1. Only the DeepCRISPR [10] model could not be retrained due to the unavailability of source code and data for pretraining. As the published model was already trained with whole DeepCRISPR dataset, the model actually had an advantage in comparison to other models. All the models were evaluated with the test set and compared with our best performing RNN, LSTM, GRU and ELECTRA models. The results are shown in Figure 4. The underlying values are reported in Appendix C.

**Fig. 4.**
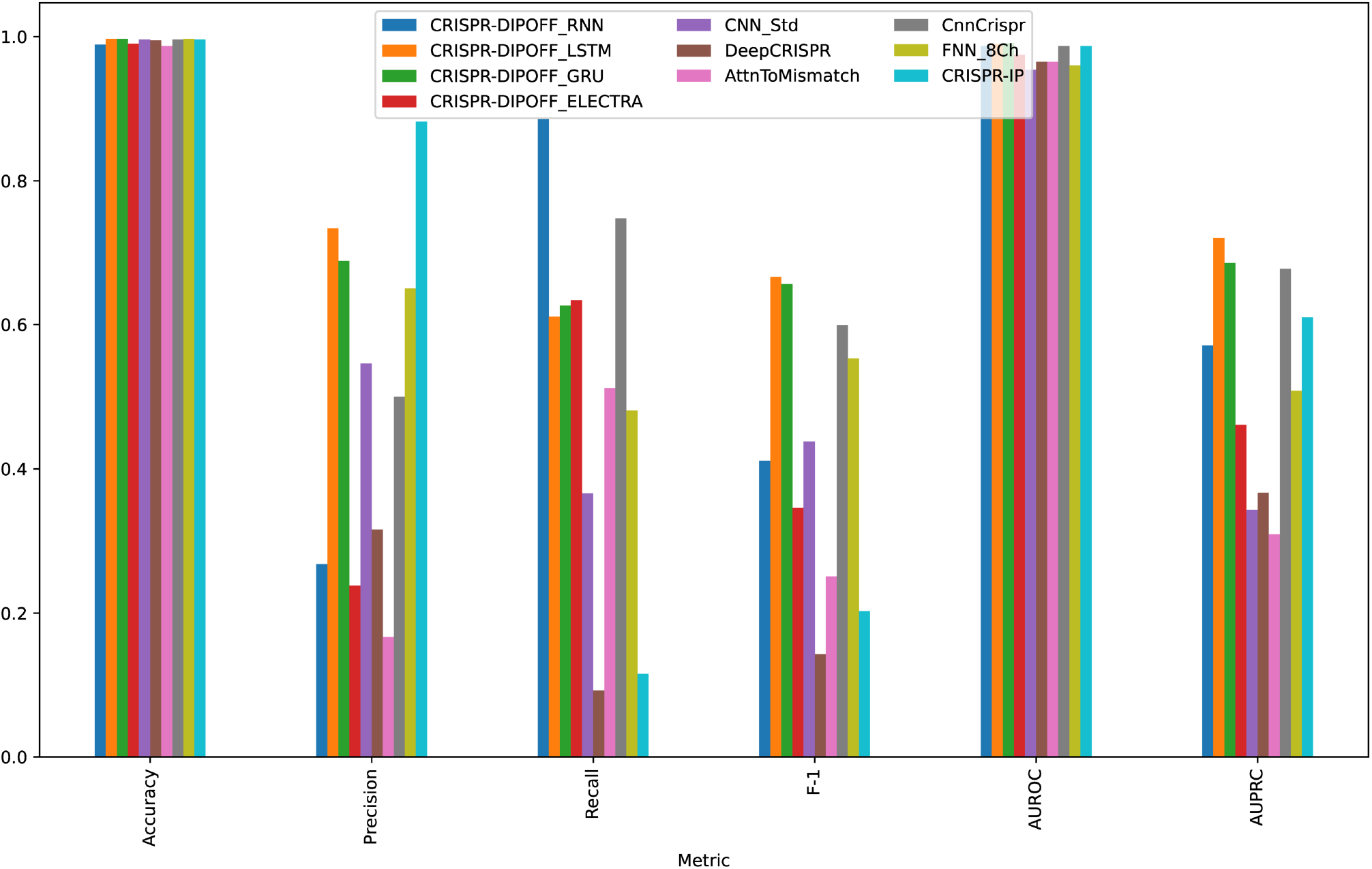
Comparison of different models based on different performance metrics. The accuracy and AUROC score are quite similar for all the models. Other metrics have varied a lot. Our LSTM model achieved the most balanced score with highest F-1 and AUPRC score.

Both our LSTM and GRU models outperformed all other models in terms of AUPRC score. Although our RNN model showed a slight decrease in performance on the test set, it still performed better than some of the previous models. Contrary to our expectation, the finetuned ELECTRA model did not yield the desired results and turned out to be the worst performing model among the four of our models. Among the baseline models, CnnCrispr demonstrated the most balanced performance, followed by CRISPR-IP. Other baseline models, such as CNN Std, DeepCRISPR, AttnToMismtach CNN, FNN 8Ch, demonstrated varying levels of performance.

Figure 4 shows detailed comparison among the models in terms of each performance metrics. As the dataset is highly imbalanced towards the negative class, it is expected that accuracy would be high even if a model is bad at predicting the positive class. Hence we observe that all the models have quite similar accuracy. The highest accuracy is 0.997 achieved by LSTM, GRU and the FFN 8Ch model. We can observe the same trend for AUROC score too. Here the best score 0f 0.991 is achieved by our GRU model. We observe varying scores across the other performance metrics. There is a clear precision recall trade-off among the models. For example, the CRISPR IP model has the highest precision (0.882) but the 2nd lowest recall. On the other hand our RNN model has the highest recall (0.886) but quite low precision. This trade-off is more prevalent in the baseline models and is evident in F-1 scores which is the harmonic mean of precision and recall. Both our LSTM and GRU models have relatively balanced precision and recall with the precision being slightly higher. Hence these two models have the highest F1 score of 0.667 and 0.656 respectively. The other performance metric is AUPRC which is considered a better indicator of performance for highly imbalanced dataset. The highest AUPRC score (0.721) was achieved by our LSTM model and it is followed by our GRU model. The LSTM model has achieved 6.34% increase in AUPRC score and 11.35% increase in F-1 score as compared to CnnCrispr which is currently the best baseline model. Overall, the recurrent neural network models, particularly LSTM, and GRU, significantly outperformed the baselines. This indicates the superiority of sequence based models and the potential of hyperparameter-tuned models for improved performance in CRISPR Off-Target prediction. Another important observation is the simplicity of the high-performing models. Further elaborations along this line are done in Section 4.1.

## Discussions

### Observation of Results

Some interesting and insightful observations may be derived from the results and analysis conducted in our experiments. Firstly, the ‘intelligent’ search mechanism for hyperparameter tuning through the use of genetic algorithms has yielded models with superior performance, despite the fact that the recurrent network-based models obtained from this technique have a very simple architecture. This observation highlights two important insights. One is the emphasis on the criticality of proper hyperparameter tuning in achieving superior performance. By carefully selecting and tuning the hyperparameters, we apparently were able to unlock the full potential of the RNN type models and surpass the performance of the more intricate and complex baseline models. The other insight aligns with the principle of Occam’s Razor, which suggests that simpler models are often more likely to be correct among competing hypotheses. In the context of our study, this implies that a well-optimized and properly tuned simpler RNN based architecture can effectively capture the underlying patterns and dependencies in CRISPR Cas-9 Off-target prediction better than more complex models.

Secondly, among the different encoding schemes used, the 4-channel encoding approach has consistently produced the best results for the RNN, LSTM, and GRU models. This finding suggests that the addition of the directional feature in 5-channel encoding may have interfered with the generalization capability of the recurrent neural network models. Alternatively, it is possible that the specific mismatch patterns themselves, rather than the direction of the mismatches, play a more significant role in determining off-target prediction. Further analysis is needed to better understand the underlying mechanisms driving the performance variations observed across different encoding schemes.

Additionally, the finetuned ELECTRA model did not yield the expected output, which can be attributed to the way we pretrained the model. Instead of training the model with complete human genome, which would have required significant time and resources, we followed the pretraining approach of DeepCRISPR by selecting samples generated by Cas-OFFinder [3]. It resulted in much smaller pretraining dataset which was manageable within our limited computational resources. However, this limited pretraining dataset may have hindered the generalizability of the model. The overlapping nature of tri-mers during training could also have led the model to easily predict a masked token based on the previous or next token, potentially resulting in earlier convergence and limited generalization. To address these limitations, a more appropriate approach would involve training the ELECTRA model with the entire human genome sequence. This would not only enhance its application for Off-Target prediction but also make it valuable for other tasks related to human genome analysis.

We note however that, despite its below par performance with respect to some of the baselines and other models in our CRISPR-DIPOFF suite, we find it worth discussing for the following reasons. Firstly, Large Language Models (LLMs) have the potential to propel the advancement of bioinformatics, similar to their impact on NLP. Secondly, despite pretraining ELECTRA in our resource constrained setting, it showed the indication of possible performance improvement. This suggests that properly pretrained ELECTA or similar LLMs could be Swiss army knife for any biological sequence related prediction tasks. Finally, the interpretations of the finetuned ELECTRA model acts as the second independent (computational) validation for our interesting observation obtained from LSTM mode’s interpretation (as discussed in a forthcoming sections).

### Model Interpretation

We have applied integrated gradients [51], an axiomatic interpretation technique, for explaining the predictions of our deep learning models. This method provides better interpretation over other methods like plain gradients, LIME (Local Interpretable Model-agnostic Explanations) [45], DeepLift [48] Layer-wise Relevance Propagation [37] etc. With Integrated Gradients, we can quantify the impact of each feature by computing the accumulated gradients along the integration path from a baseline input to the actual input. The “baseline” is the reference input, generally an input containing all zeros, that is expected to have no effect on the classification and the “actual input” is the specific data point we want to interpret. Layers and even neurons can also be interpreted with this method.

### Interpretation of the LSTM Model

In our study, the best performing model is a LSTM model which is trained on the 4-channel encoded input. This model’s architecture is shown in Figure 5. The model has a bidirectional LSTM layer followed by two hidden layers and the output layer. The hidden layers have 512 and 256 neurons respectively.

**Fig. 5.**
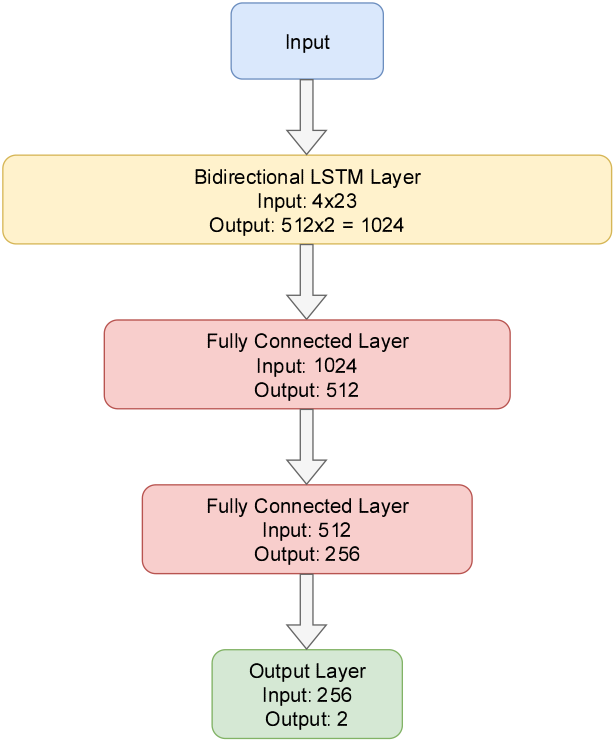
Architecture of Best Performing LSTM Model. The model has a bidirectional LSTM followed by two hidden layers and an output layer.

#### Feature Importance

We calculated attribution scores for all the test samples using Integrated Gradients with respect to positive class. For each sample, we normalized the attribution scores for all the features present in that sample. After that we averaged the score of each feature across all the samples. Based on this score, top 15 contributing features for positive, negative and overall prediction of the model are shown in Figure 6. Due to the higher ratio of negative samples, the most contributing features for negative and overall prediction are almost the same. We have drawn a few major observations from the importance scores as follows.

**Fig. 6.**
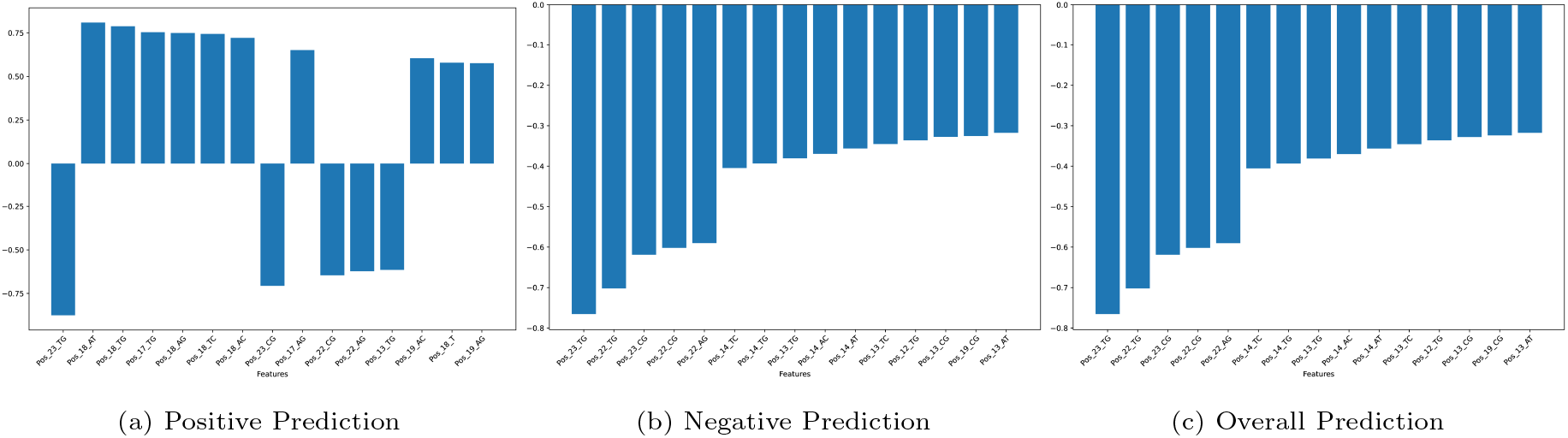
Top 15 features contributing most to (a) positive, (b) negative and (c) overall prediction with respect to positive class.

- **The effects of Mismatch**: All the top contributing features are related to mismatches. This is intuitive as mismatch is the main reason of Off-Target. After ranking the features based on attribution score, we found that 38 out of top 50 features for positive prediction are mismatch type features. In case of both negative and overall prediction, top 44 out of 50 features are mismatch type features.
- **Different Regions in sgRNA**: The 23-mer sequence can be divided in three main regions: PAM (position 21-23), seed or PAM proximal (5 to 12 nucleotides upstream or near the PAM sequence) and PAM distal (rest of the nucleotides) [56]. The 21st nucleotide or the first nucleotide of PAM is ‘N’ which is similar to a wildcard for nucleotides and hence we can observe that none of our top contributing features indicate mismatch at the 21st nucleotide. The other two nucleotide of PAM contains ‘GG’ for Cas-9 and we observe that mismatch with ‘G’ at these two position heavily contribute in both positive and negative predictions. The features contributing to the positive predictions indicate that Positions 17, 18 and 19 have strong positive correlation with the positive class. For negative and overall prediction, all the top ranked features demonstrate negative correlation and the top most features are mismatches at the PAM region. The other features correspond to mismatches at the seed regions specially at positions 12 to 14.
- **Sub-regions in the Seed Region**: The total number of positively correlated mismatch features for positive prediction are 45 and among them half are at the PAM proximal region (positions 16 to 20) and the rest are at the PAM distal region (positions 1 to 10). The mismatches at positions 11 to 15 have negative correlation. On the other hand, for negative and overall prediction only 12 mismatch type features have positive correlation and 11 of them are at the PAM proximal region (positions 17 to 20) and the other one is at the PAM distal region. Previous studies have identified that mismatches close to PAM, specially in the so called seed region, decrease the likelihood of Off-Target effects and mismatches at the PAM distal region are sometimes tolerated during sgRNA-DNA binding which results in Off-Target effect [2, 15, 23]. The observation of features for positive prediction aligns with this established hypothesis at least partially. But our model’s interpretation further suggests quite strongly that probably there are two separate portions in the seed regions. One portion is closer to the PAM (positions 16/17 to 20) and the other is adjacent to the PAM distal region (positions 11 to 15/16). According to Zheng at. al. [61] there is a core region inside the seed region and mismatch therein strongly affects the cleavage activity of CRISPR. The 2nd portion of the seed region detected in our study closely overlaps with the core region. It is possible that the deep learning model has learned to distinguish certain features in a manner that somewhat extends the established hypothesis. It also highlights the complexity of interpreting the inner workings of deep learning models and the need for further investigation to reconcile these findings with existing biological knowledge.
- **Mismatch containing ‘TG’ and ‘CG’**: A closer observation of the top ranked features reveals that mismatches are dominated by ‘TG’ and ‘CG’ type substitutions. This was also reported in DeepCRISPR [10] as a distinguishable property.

#### Layer Importance

Layer attributions calculated with integrated gradients was used to interpret layer importance. Layer attributions provide us with an understanding of the significance of all the neurons contributing to the output of a specific layer. For each test sample, attribution score of each neuron of each layer was calculated. These scores were averaged across positive, negative and all samples. In our LSTM model, there are two hidden layers. The layer immediately after the Bi-Directional LSTM has 512 neurons and the next layer has 256 neurons. The top 15 contributing neurons on positive, negative and overall predictions are shown in Figure 7.

**Fig. 7.**
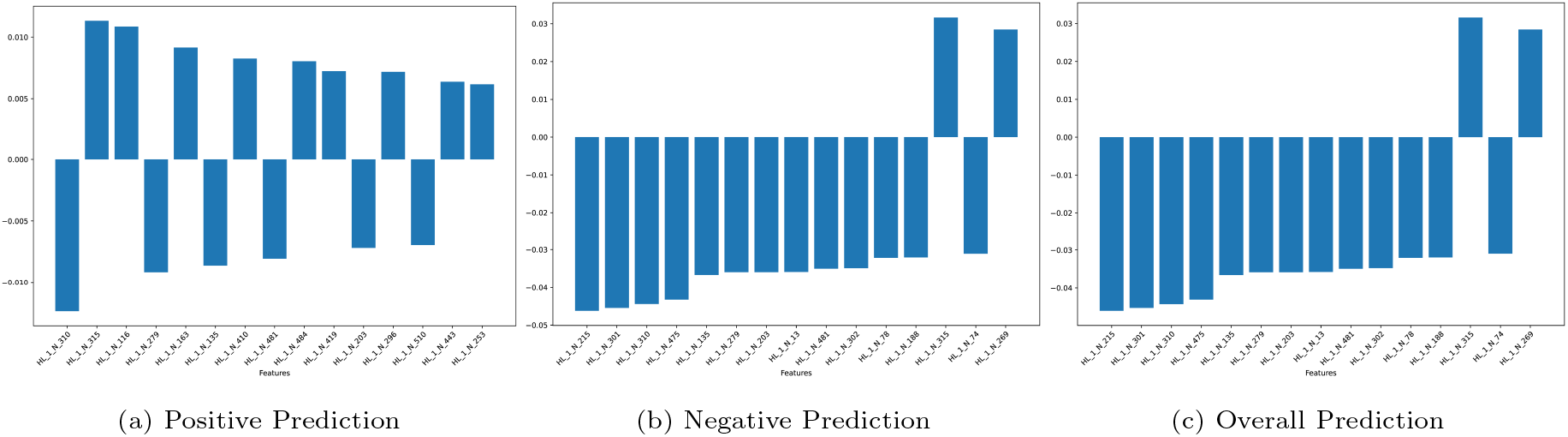
Top 15 neurons contributing most to (a) positive, (b) negative and (c) overall prediction with respect to positive class.

It is noticeable that all the neurons that have higher attribution scores are from the 1st hidden layer. In fact, only 16 out of top 100 neurons in terms of overall prediction are from the 2nd hidden layer. This clearly indicates that the 1st hidden layer has more leverage on the final model output. Also, the output of the LSTM layer is directly fed to the 1st hidden layer. Hence it is possible that the 1st hidden layer is being able to capture more generalized context. The dominance of 1st hidden layer over the output is clearly visible from the heatmaps shown in Figure 8. The heatmap of the 1st hidden layer is filled with reddish color which indicates more active neurons in this layer. On the other hand, many of the neurons of the 2nd hidden layers are inactive or minimally active which is denoted by the dark patches.

**Fig. 8.**
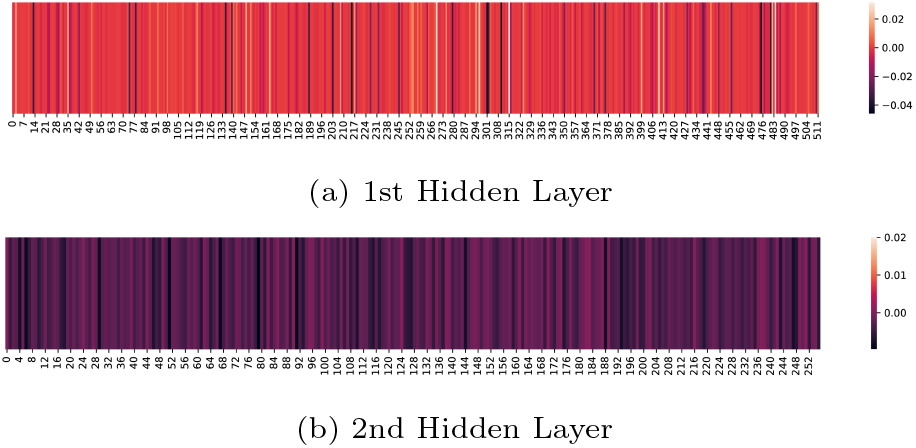
Activation heatmap of neurons of (a) 1st and (b) 2nd hidden layer

### Interpretation of the ELECTRA Model

The ELECTRA model, finetuned for Off-Target prediction, did not perform as expected. Among the different ELECTRA models, the pretrained model on tri-mer encoded tokens and finetuned with two additional layers performed better than others. Despite not meeting our initial performance expectations, we have attempted to interpret the model in a limited capacity. Given the complexity and size of the model, which comprises approximately 3.2 million parameters, interpreting its inner workings becomes an arduous task. To overcome this challenge, we focused our interpretation efforts on the embedding layers of the finetuned ELECTRA model. The ELECTRA model consists of three specific embedding layers for token, token type, and position embedding. We employed the integrated gradients method to calculate attribution scores for these layers. The input tokens used in our experiments are tri-mer encoded sgRNA and DNA sequences, separated by [SEP] tokens. There is also [CLS] token at the start and a [SEP] token at the end. In order to align with the maximum sequence length, last three empty positions were filled with [PAD] tokens. Integrated gradients require a baseline sample to compute the gradient along the path. Generally an input vector containing all zeros is used for that purpose. In case of ELECTRA, the baseline sample has bee prepared with [PAD] tokens.

We computed the average attribution scores for each tokens across positive, negative, and overall predictions on the finetuned model. To observe the change in embedding layer we also computed attribution scores for overall predictions on pretrained model (i.e., the model before finetuning). The token importances are illustrated in Figure 9 which clearly demonstrates that the tokens associated with the DNA sequence exhibit significantly higher attribution scores compared to the sgRNA tokens. This observation is well-aligned with the understanding that the sgRNA sequence is engineered, while the target DNA sequence is responsible for introducing mismatches. In practical scenarios, a single sgRNA can potentially interact with numerous off-target DNA sites, and the final determination of off-target effects depends on the mismatches introduced by the DNA sequences. It appears that the embedding layers of the finetuned ELECTRA model have successfully captured this relationship from the training data used during finetuning. The comparison of attribution before and after the finetuning process, illustrates how the embedding layer has changed after the finetuning process and has been aware of the biological significance of the specific task of Off-Target prediction.

**Fig. 9.**
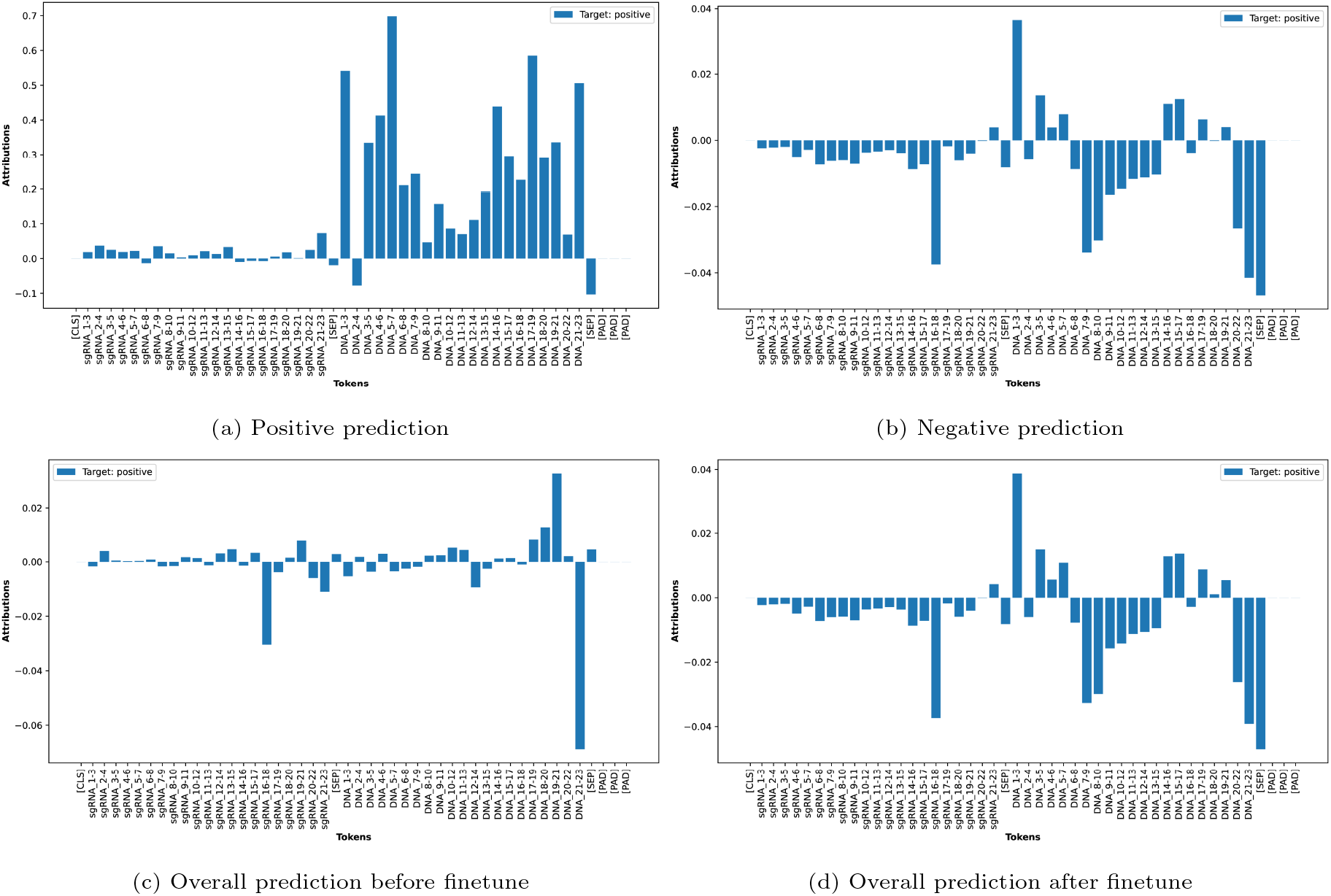
Attribution scores at embedding layers for all the input tokens of finetuned ELECTRA model for (a) positive, (b) negative and (c) overall prediction before finetune and d) overall prediction after finetune with respect to positive class.

Similar to the interpretation of our LSTM model, we observe that the attribution scores for negative and overall prediction are almost the same. The attribution scores for overall predictions show some sort of four distinguishable regions in the DNA tokens. The first region is related to PAM and it affects the final prediction negatively. There are two regions with contiguous positive scores. One of them is in the seed region (position 15 to 20) and the other one is in PAM distal region (position 1 to 6). The region in the middle to these two regions contain contiguous negative attributions. This is consistent with our findings in LSTM model’s interpretation, though the regions does not match exactly nucleotide by nucleotide. This further strengthens our observation that there are two sub-regions in the seed regions and one of them might be tolerable for mismatches. Further investigation is required to validate this observation. We anticipate that improving the performance with large scale pretraining of ELECTRA model could unlock more complex biological relationship and deeper understanding of Off-Target effects.

## Conclusion

CRISPR Cas-9 has shown great promises in the field of gene editing for the past few years but the possibility of Off-Target effects still remain as the monumental challenge. The current state-of-the-art deep learning models suffer from precision-recall trade-off and due to the lack of interpretations, these models are still black boxes to the end users. In this study, we have successfully developed interpretable deep learning models capable of predicting Off-Target sites with high efficacy surpassing the previous studies using only sequence data. Through the interpretation of our models we seem to have unraveled the existence of sub-regions within the seed region. This can be seen as an extension/elaboration of the current hypothesis and it calls for further biological and computational research. Future computational endeavours should also focus on pretraining ELECTRA or similar models, particularly on whole genome sequence, which could be used for both On-Target and Off-Target predictions at the same time and even for any other predictive tasks requiring genomic contexts.

## Appendix

### A. Model Selection with Genetic Algorithm

In order to perform model selection, we ran plain Genetic Algorithm (GA) and Elitist Genetic Algorithm (EGA) separately for 20 generations and each generation had 20 models. We ran separate experiments for 4-channel and 5-channel encoding scheme. Including the initial population, we trained a total of 420 models in each experiment. The same experiment was run independently for RNN, LSTM and GRU type recurrent networks. As each of these models work a bit differently, it was not conducive to perform crossover among different types of recurrent networks. Hence we could not include model type (RNN, LSTM or GRU) as another hyperparameter for our tuning algorithm. Notably, instead of checking 72000 (or even more) possible models for each type of recurrent networks, we only, but intelligently, explored 420 thereof. We ranked the models based on their AUPRC score on the validation set.

The results of plain genetic algorithm on 4-channel and 5-channel encoded input are shown in Table 1 and Table 2 respectively. We have shown results for top 3 models for each recurrent network types. According to the result, all the models have performed quite well on the validation set compared to the baselines in terms of AUPRC score. But it is interesting to observe that most of the models were already found on iteration 0, 1 or 2. Later generations did not improve the performance. This indicated that the best performing models from initial generation were not preserved in the subsequent generations and some useful parameter values might have got lost. Hence we used Elitist Genetic Algorithm to preserve the best performing models. We also draw an interesting observation that most of the best performing models have hidden size 512.

The experiments using Elitist Genetic Algorithm (EGA) was similar to the plain Genetic Algorithm approach except that we preserved four best performing models of the current generation for the next generation. The results of elitist genetic algorithm on 4-channel and 5-channel encoded input are shown in Table 3 and Table 4 respectively. The tables reveal that the models have gradually improved over subsequent iterations. The performance in terms of AUROC is quite well on 4-channel encoded input. Surprisingly, in both GA and EGA, performance of 5-channel encoded data is worse. This observation suggests that the mismatch itself might be the primary factor influencing Off-Target prediction, and incorporating the direction of the mismatch as an engineered feature may have introduced unnecessary noise or complexity to the models.

**Table 1.** Parameters and Results of Best RNN, LSTM, and GRU Models Obtained from Genetic Algorithm with 4-Channel Encoded Input. AUPRC score has been calculated on Validation set.

| Model Type | Iteration | Hidden Size | LSTM Layers | Bi LSTM | Hidden Layers | Dropout Probability | Batch Size | Epochs | Learning Rate | AUPRC |
| --- | --- | --- | --- | --- | --- | --- | --- | --- | --- | --- |
| RNN | 3 | 512 | 1 | FALSE | 2 | 0.4 | 256 | 30 | 0.00050 | 0.7080 |
|  | 2 | 512 | 2 | TRUE | 2 | 0.1 | 64 | 50 | 0.00005 | 0.6914 |
|  | 2 | 512 | 1 | FALSE | 2 | 0.2 | 256 | 20 | 0.00010 | 0.6578 |
| LSTM | 1 | 512 | 2 | FALSE | 2 | 0.2 | 64 | 20 | 0.00005 | 0.7324 |
|  | 0 | 64 | 1 | TRUE | 4 | 0.15 | 256 | 30 | 0.00100 | 0.7203 |
|  | 0 | 512 | 2 | TRUE | 2 | 0.1 | 64 | 50 | 0.00005 | 0.7120 |
| GRU | 0 | 128 | 2 | TRUE | 0 | 0.1 | 64 | 30 | 0.00050 | 0.7427 |
|  | 1 | 512 | 2 | FALSE | 2 | 0.25 | 64 | 20 | 0.00005 | 0.7410 |
|  | 1 | 512 | 2 | FALSE | 2 | 0.2 | 64 | 20 | 0.00005 | 0.7016 |

**Table 2.** Parameters and Results of Best RNN, LSTM, and GRU Models Obtained from Genetic Algorithm with 5-Channel Encoded Input. AUPRC score has been calculated on Validation set.

| Model Type | Iteration | Hidden Size | LSTM Layers | Bi LSTM | Hidden Layers | Dropout Probability | Batch Size | Epochs | Learning Rate | AUPRC |
| --- | --- | --- | --- | --- | --- | --- | --- | --- | --- | --- |
| RNN | 3 | 512 | 1 | FALSE | 2 | 0.1 | 32 | 90 | 0.00010 | 0.7135 |
|  | 0 | 512 | 1 | FALSE | 2 | 0.15 | 256 | 30 | 0.00050 | 0.7045 |
|  | 1 | 128 | 2 | FALSE | 0 | 0.1 | 256 | 20 | 0.00100 | 0.6983 |
| LSTM | 1 | 64 | 2 | TRUE | 4 | 0.5 | 256 | 30 | 0.00500 | 0.7157 |
|  | 0 | 64 | 1 | TRUE | 4 | 0.15 | 256 | 30 | 0.00100 | 0.6890 |
|  | 2 | 512 | 1 | FALSE | 2 | 0.4 | 64 | 50 | 0.00050 | 0.6885 |
| GRU | 2 | 512 | 1 | FALSE | 2 | 0.4 | 64 | 50 | 0.00050 | 0.6885 |
|  | 3 | 64 | 2 | FALSE | 1 | 0.1 | 256 | 30 | 0.00100 | 0.6872 |
|  | 5 | 256 | 1 | TRUE | 2 | 0.45 | 32 | 50 | 0.00010 | 0.6860 |

**Table 3.** Parameters and Results of Best RNN, LSTM, and GRU Models Obtained from Elitist Genetic Algorithm with 4-Channel Encoded Input. AUPRC score has been calculated on Validation set.

| Model Type | Iteration | Hidden Size | LSTM Layers | Bi LSTM | Hidden Layers | Dropout Probability | Batch Size | Epochs | Learning Rate | AUPRC |
| --- | --- | --- | --- | --- | --- | --- | --- | --- | --- | --- |
| RNN | 4 | 256 | 2 | TRUE | 0 | 0.4 | 256 | 60 | 0.00100 | 0.7643 |
|  | 3 | 128 | 1 | FALSE | 4 | 0.2 | 128 | 30 | 0.00100 | 0.7417 |
|  | 9 | 128 | 1 | TRUE | 4 | 0.15 | 128 | 30 | 0.00100 | 0.7284 |
| LSTM | 1 | 512 | 1 | TRUE | 2 | 0.4 | 64 | 50 | 0.00010 | 0.7403 |
|  | 7 | 512 | 1 | TRUE | 2 | 0.1 | 256 | 80 | 0.00005 | 0.6998 |
|  | 1 | 64 | 1 | TRUE | 4 | 0.15 | 256 | 30 | 0.00100 | 0.6980 |
| GRU | 1 | 512 | 1 | FALSE | 2 | 0.4 | 64 | 50 | 0.00010 | 0.7403 |
|  | 1 | 128 | 1 | FALSE | 4 | 0.2 | 64 | 50 | 0.00010 | 0.7168 |
|  | 1 | 512 | 2 | FALSE | 2 | 0.1 | 64 | 50 | 0.00005 | 0.7079 |

**Table 4.** Parameters and Results of Best RNN, LSTM, and GRU Models Obtained from Elitist Genetic Algorithm with 5-Channel Encoded Input. AUPRC score has been calculated on Validation set.

| Model Type | Iteration | Hidden Size | LSTM Layers | Bi LSTM | Hidden Layers | Dropout Probability | Batch Size | Epochs | Learning Rate | AUPRC |
| --- | --- | --- | --- | --- | --- | --- | --- | --- | --- | --- |
| RNN | 1 | 512 | 1 | FALSE | 2 | 0.15 | 256 | 30 | 0.00050 | 0.7058 |
|  | 3 | 512 | 2 | FALSE | 2 | 0.1 | 256 | 80 | 0.00100 | 0.6951 |
|  | 6 | 512 | 2 | FALSE | 2 | 0.2 | 256 | 80 | 0.00050 | 0.6754 |
| LSTM | 3 | 512 | 2 | FALSE | 2 | 0.1 | 256 | 80 | 0.00100 | 0.6951 |
|  | 6 | 512 | 2 | FALSE | 2 | 0.2 | 256 | 80 | 0.00050 | 0.6754 |
|  | 7 | 512 | 1 | TRUE | 2 | 0.1 | 256 | 80 | 0.00005 | 0.6266 |
| GRU | 4 | 512 | 2 | FALSE | 2 | 0.1 | 256 | 80 | 0.00100 | 0.6951 |
|  | 6 | 512 | 2 | FALSE | 2 | 0.2 | 256 | 80 | 0.00050 | 0.6754 |
|  | 7 | 512 | 1 | TRUE | 2 | 0.1 | 256 | 80 | 0.00005 | 0.6266 |

### B. Finutuned ELECTRA Model

During the finetuning phase of the ELECTRA models, we employed a strategic approach to enhance its performance. This involved incorporating additional layers, ranging from 1 to 3, including the output layer, specifically designed for classification purposes. By introducing these extra layers, we aimed to capture more intricate patterns and improve the model’s ability to make accurate predictions. To finetune the ELECTRA models, we utilized the gradual unfreezing strategy [22]. This technique allows for a controlled update of the model’s weights, starting with the output and additional layers and progressively unfreezing the encoder layers closest to it. In our experiments, we conducted separate finetuning procedures on the Tiny models that were pretrained using both Tri-Mer Encoded (TME) and Byte Pair Encoded (BPE) input tokens. The results of these experiments, as shown in Table 5 and Table 6 for the TME and BPE models, respectively, shed light on the impact of these encoding approaches on the model’s performance. Notably, we observed a significant performance advantage for the TME models compared to the BPE models. Among the TME models, the one that yielded the best results incorporated two additional layers, an additional hidden layer and an output layer, following the ELECTRA encoder layers.

Though the overall result does not outperform the baseline models and the RNN based models, we observe that gradual unfreezing improved the performance of the models. Gradual unfreezing provides better results because it helps to mitigate the problem of catastrophic forgetting. Catastrophic forgetting is a phenomenon that occurs when a machine learning model is finetuned on a new task, and it loses its ability to perform well on the original task. While finetuning the ELECTRA model, we are essentially updating the weights of the model to better fit the new task. However, if we update all of the weights at once, it is possible that the model will forget how to perform the original task which is capturing the general context of the sequences. Gradual unfreezing helps to mitigate this problem by unfreezing the layers of the model from the last layer to the first layer. The last layer of the model contains the least general knowledge, so it is the least likely to be affected by finetuning on a new task.

**Table 5.** Results of finetuned ELECTRA models pretrained with Tri-Mer Encoded (TME) input tokens. The result shows how the performance improved over the gradual unfreezing of ELECTRA layers.

| Model Type | Unfrozen Layers | Trainable ELECTRA Parameters | Accuracy | Precision | Recall | F1-Score | AUROC | AUPRC |
| --- | --- | --- | --- | --- | --- | --- | --- | --- |
| TME with 1 FCL | None | 0 | 0.7888 | 0.0174 | 0.8702 | 0.0340 | 0.8772 | 0.0261 |
|  | Layer 6 | 527104 | 0.8237 | 0.0219 | 0.9237 | 0.0429 | 0.9377 | 0.1035 |
|  | Layer 5 to 6 | 1054208 | 0.8230 | 0.0219 | 0.9237 | 0.0427 | 0.9452 | 0.1169 |
|  | Layer 4 to 6 | 1581312 | 0.8315 | 0.0240 | **0.9695** | 0.0469 | 0.9585 | 0.1398 |
|  | Layer 3 to 6 | 2108416 | 0.9394 | 0.0597 | 0.8931 | 0.1120 | 0.9775 | 0.3407 |
|  | Layer 2 to 6 | 2635520 | 0.9782 | 0.1365 | 0.7710 | 0.2319 | 0.9760 | 0.4081 |
|  | Layer 1 to 6 | 3162624 | 0.9897 | 0.2409 | 0.6565 | 0.3525 | **0.9795** | 0.4236 |
|  | All Layers | 3194112 | **0.9918** | **0.2842** | 0.6031 | **0.3863** | 0.9542 | 0.4288 |
| TME with 2 FCL | None | 0 | 0.8078 | 0.0197 | 0.9008 | 0.0385 | 0.9264 | 0.0657 |
|  | Layer 6 | 527104 | 0.8077 | 0.0208 | 0.9542 | 0.0407 | 0.9369 | 0.1009 |
|  | Layer 5 to 6 | 1054208 | 0.8244 | 0.0229 | 0.9618 | 0.0447 | 0.9479 | 0.1078 |
|  | Layer 4 to 6 | 1581312 | 0.8243 | 0.0231 | **0.9695** | 0.0450 | 0.9572 | 0.1161 |
|  | Layer 3 to 6 | 2108416 | 0.9380 | 0.0576 | 0.8779 | 0.1080 | 0.9752 | 0.2846 |
|  | Layer 2 to 6 | 2635520 | 0.9772 | 0.1271 | 0.7405 | 0.2170 | 0.9775 | 0.3596 |
|  | Layer 1 to 6 | 3162624 | 0.9857 | 0.1848 | 0.6870 | 0.2913 | 0.9742 | 0.3951 |
|  | All Layers | 3194112 | 0.9898 | 0.2378 | 0.6336 | 0.3458 | 0.9752 | **0.4613** |
| TME with 3 FCL | None | 0 | 0.8256 | 0.0220 | 0.9160 | 0.0430 | 0.9385 | 0.0777 |
|  | Layer 6 | 527104 | 0.8127 | 0.0208 | 0.9313 | 0.0408 | 0.9363 | 0.0980 |
|  | Layer 5 to 6 | 1054208 | 0.8024 | 0.0206 | **0.9695** | 0.0402 | 0.9435 | 0.1095 |
|  | Layer 4 to 6 | 1581312 | 0.7963 | 0.0203 | 0.9847 | 0.0397 | 0.9576 | 0.1129 |
|  | Layer 3 to 6 | 2108416 | 0.9464 | 0.0660 | 0.8779 | 0.1228 | 0.9757 | 0.2432 |
|  | Layer 2 to 6 | 2635520 | 0.9758 | 0.1225 | 0.7557 | 0.2109 | 0.9756 | 0.2317 |
|  | Layer 1 to 6 | 3162624 | 0.9875 | 0.2033 | 0.6565 | 0.3105 | 0.9688 | 0.3995 |
|  | All Layers | 3194112 | 0.9881 | 0.2118 | 0.6565 | 0.3203 | 0.9746 | 0.3896 |

**Table 6.** Results of finetuned ELECTRA models pretrained with Byte Pair Encoded (BPE) input tokens. The result shows how the performance improved over the gradual unfreezing of ELECTA layers.

| Model Type | Unfrozen Layers | Trainable ELECTRA Parameters | Accuracy | Precision | Recall | F1-Score | AUROC | AUPRC |
| --- | --- | --- | --- | --- | --- | --- | --- | --- |
| BPE with 1 FCL | None | 0 | 0.6538 | 0.0106 | 0.8702 | 0.0210 | 0.8174 | 0.0186 |
|  | Layer 6 | 527104 | 0.7193 | 0.0130 | 0.8626 | 0.0256 | 0.8767 | 0.0469 |
|  | Layer 5 to 6 | 1054208 | 0.7463 | 0.0145 | 0.8702 | 0.0285 | 0.8892 | 0.0564 |
|  | Layer 4 to 6 | 1581312 | 0.8089 | 0.0183 | 0.8321 | 0.0359 | 0.9053 | 0.0774 |
|  | Layer 3 to 6 | 2108416 | 0.8714 | 0.0257 | 0.7863 | 0.0497 | 0.9234 | 0.0990 |
|  | Layer 2 to 6 | 2635520 | 0.9621 | 0.0620 | 0.5573 | 0.1116 | 0.9282 | 0.1716 |
|  | Layer 1 to 6 | 3162624 | 0.9832 | 0.1203 | 0.4656 | 0.1912 | 0.9236 | 0.1681 |
|  | All Layers | 3266816 | 0.9879 | 0.1653 | 0.4504 | 0.2418 | 0.9213 | 0.2321 |
| BPE with 2 FCL | None | 0 | 0.7275 | 0.0123 | 0.7939 | 0.0243 | 0.8545 | 0.0288 |
|  | Layer 6 | 527104 | 0.7377 | 0.0139 | 0.8626 | 0.0273 | 0.8809 | 0.0559 |
|  | Layer 5 to 6 | 1054208 | 0.7583 | 0.0156 | 0.8931 | 0.0306 | 0.8950 | 0.0534 |
|  | Layer 4 to 6 | 1581312 | 0.8244 | 0.0196 | 0.8168 | 0.0382 | 0.9098 | 0.0696 |
|  | Layer 3 to 6 | 2108416 | 0.8890 | 0.0280 | 0.7405 | 0.0539 | 0.9217 | 0.1130 |
|  | Layer 2 to 6 | 2635520 | 0.9610 | 0.0603 | 0.5573 | 0.1088 | 0.9260 | 0.1606 |
|  | Layer 1 to 6 | 3162624 | 0.9784 | 0.0945 | 0.4733 | 0.1576 | 0.9251 | 0.1725 |
|  | All Layers | 3266816 | 0.9889 | 0.1745 | 0.4275 | 0.2478 | 0.9220 | 0.1788 |
| BPE with 3 FCL | None | 0 | 0.7705 | 0.0142 | 0.7710 | 0.0279 | 0.8671 | 0.0346 |
|  | Layer 6 | 527104 | 0.7058 | 0.0130 | 0.9084 | 0.0257 | 0.8739 | 0.0482 |
|  | Layer 5 to 6 | 1054208 | 0.7350 | 0.0139 | 0.8702 | 0.0273 | 0.8849 | 0.0586 |
|  | Layer 4 to 6 | 1581312 | 0.7871 | 0.0166 | 0.8397 | 0.0326 | 0.9053 | 0.0749 |
|  | Layer 3 to 6 | 2108416 | 0.8908 | 0.0290 | 0.7557 | 0.0558 | 0.9205 | 0.0985 |
|  | Layer 2 to 6 | 2635520 | 0.9597 | 0.0555 | 0.5267 | 0.1004 | 0.9224 | 0.1360 |
|  | Layer 1 to 6 | 3162624 | 0.9731 | 0.0830 | 0.5267 | 0.1435 | 0.9209 | 0.1141 |
|  | All Layers | 3266816 | 0.9864 | 0.1439 | 0.4427 | 0.2172 | 0.9156 | 0.1969 |

### C. Comparison of Performance

Comparison of our models with previous studies is shown in Table 7.

**Table 7.** Comparison of results of our study with baseline studies. The models of our study have the prefix “CRISPR-DIPOFF”. Our LSTM model have outperformed the models from previous studies by a fair margin.

| Model Name | Accuracy | Precision | Recall | F1-Score | AUROC | AUPRC |
| --- | --- | --- | --- | --- | --- | --- |
| CRISPR-DIPOFF_RNN | 0.989 | 0.268 | **0.886** | 0.411 | 0.987 | 0.571 |
| CRISPR-DIPOFF_LSTM | **0.997** | 0.734 | 0.611 | **0.667** | 0.990 | **0.721** |
| CRISPR-DIPOFF_GRU | **0.997** | 0.689 | 0.626 | 0.656 | **0.991** | 0.686 |
| CRISPR-DIPOFF_ELECTRA | 0.990 | 0.238 | 0.634 | 0.346 | 0.975 | 0.461 |
| CNN_Std [33] | 0.996 | 0.546 | 0.366 | 0.438 | 0.954 | 0.343 |
| DeepCRISPR [10] | 0.995 | 0.316 | 0.092 | 0.142 | 0.965 | 0.367 |
| AttnToMismtach_CNN [35] | 0.987 | 0.166 | 0.512 | 0.251 | 0.965 | 0.309 |
| CnnCrispr [36] | 0.996 | 0.500 | 0.748 | 0.599 | 0.987 | 0.678 |
| FNN_8Ch [5] | **0.997** | 0.650 | 0.481 | 0.553 | 0.960 | 0.508 |
| CRISPR-IP [60] | 0.996 | **0.882** | 0.115 | 0.203 | 0.987 | 0.610 |

## Footnotes

1 https://github.com/tzpranto/CRISPR-DIPOFF

2 https://ftp.ensembl.org/pub/release-110/fasta/homo_sapiens/dna/

